# Chromosomal evolution and speciation inferred from chromosome-scale genome assemblies in three Cupressaceae species

**DOI:** 10.64898/2026.01.06.698058

**Authors:** Kenta Shirasawa, Yuta Aoyagi Blue, Kentaro Mishima, Hideki Hirakawa, Tomonori Hirao

**Author notes:** To whom correspondence should be addressed: Kenta Shirasawa, 2-6-7 Kazusa-Kamatari, Kisarazu, Chiba 292-0818, Japan.

## Abstract

Chromosome-scale genome assemblies in gymnosperms have lagged behind those of angiosperms, likely due to their large genomes. Coniferous tree species, which belong to the gymnosperms, are important resources for wood production in the forestry industry. To elucidate the evolution and speciation of these species and establish genome resources for breeding, we integrated draft assemblies with optical and genetic mapping to construct chromosome-scale genomes for Japanese cypress (*Chamaecyparis obtusa*, 8.7 Gb), Japanese cedar (*Cryptomeria japonica*, 9.6 Gb), and Chinese fir (*Cunninghamia lanceolata*, 13.4 Gb). Additionally, we assembled and annotated their chloroplast and mitochondrial genomes.

Comparative analysis of the nuclear genomes revealed that while synteny is largely conserved, distinct translocations and inversions occurred in chromosomes 2, 6, and 9. Notably, the significantly larger genome of *C. lanceolata* was associated with frequent tandem gene duplications rather than transposon expansion. These findings suggest that chromosomal rearrangements and segmental duplications played key roles in the divergence of these species. The genomic resources presented here including chromosome-scale sequences, gene annotations, and genetic maps will facilitate advanced conifer genetics and accelerate forest tree breeding programs.

## Introduction

The Spermatophyta, or seed plants, are broadly categorized into angiosperms and gymnosperms. Whereas angiosperms include more than 60 orders (Chase *et al*. 2016), gymnosperms currently consist of only eight orders (Christenhusz *et al*. 2011): Araucariales, Cupressales, Cycadales, Ephedrales, Ginkgoales, Gnetales, Pinales, and Welwitschiales. Although the classification of both angiosperms and gymnosperms has been debated, molecular techniques have played a significant role in resolving these arguments. Gene sequences of organelle genomes, which are conserved across species yet slightly divergent, are recognized as DNA barcodes for identifying plant species (Hebert *et al*. 2003). More recently, with great advancements in genome sequencing technologies, nuclear genome sequences at the chromosome level have been applied to the classification of angiosperms (Zhao *et al*. 2021). Even though the genome sizes of angiosperms range from 100 Mb to 100 Gb (Pellicer and Leitch 2020), sequenced plant species have been biased toward those with small genome sizes (<1 Gb) (Shirasawa *et al*. 2021). In gymnosperms, on the other hand, most species possess massive genomes, approximately 10 Gb or more (Pellicer and Leitch 2020). Therefore, the assembly of chromosome-scale genome sequences in gymnosperms has lagged behind that of angiosperms.

Among gymnosperms, the orders Pinales, Cupressales, and Araucariales are collectively known as conifers (Coniferophyta). Coniferous trees are vital resources for the forestry industry and for housing materials. In Japan, approximately 60% of the land is forested; of this, 40% comprises plantation (artificial) forests dominated by Japanese cedar (*Cryptomeria japonica*), followed by Japanese cypress (*Chamaecyparis obtusa*) (Forestry Agency 2022, Tsumura *et al*. 2007). Recently, Chinese fir (*Cunninghamia lanceolata*) has been recognized as a promising new woody resource due to its fast growth and desirable wood properties (Fujisawa 2017). However, breeding programs for coniferous trees are generally time-consuming because their generation times and harvest rotations often exceed 50 years. Genomic information would help accelerate these breeding programs by identifying genes that control traits of importance to the forestry industry. In our previous study (Shirasawa *et al*. 2024), we released draft genome sequences for *C. obtusa*, *C. japonica*, and *C. lanceolata*; however, these sequences were highly fragmented and lacked gene annotations.

Comparative analysis of genome structures using chromosome-scale assemblies can elucidate the evolution and speciation processes of species (Aoyagi Blue *et al*. 2025). Moreover, gene annotations, combined with chromosome-scale sequences, are valuable resources for identifying genes for use in breeding programs (Shirasawa *et al*. 2021). In this study, we extended the contiguity of the draft sequences for *C. obtusa*, *C. japonica*, and *C. lanceolata* using optical mapping and anchored the sequences to chromosomes using a genetic mapping strategy. Comparative genomics using the resulting chromosome-scale sequences provided insights into the evolution and speciation of these three coniferous tree species.

## Materials and methods

### Plant materials

We used the same individual trees of *C. obtusa* (tree ID: GFB00119), *C. japonica* (GFA01029), and *C. lanceolata* (GFHN00090) that were employed for the draft genome assemblies in our previous study (Shirasawa *et al*. 2024). For genetic mapping, we utilized four mapping populations for *C. obtusa*, five for *C. japonica*, and two for *C. lanceolata*.

### Optical mapping

Genomic DNA was extracted from young leaves of the three species using the Plant DNA Isolation Kit (Bionano Genomics, San Diego, CA, USA), following the Bionano Prep Plant Tissue DNA Isolation Base Protocol. The isolated genomic DNA was treated with the DLE-1 enzyme and labeled with a fluorescent dye supplied in the DLS DNA Labeling Kit (Bionano Genomics). The labeled DNA was analyzed on the Saphyr Optical Genome Mapping Instrument (Bionano Genomics). The resulting molecules were assembled and merged with the draft assembly to generate hybrid scaffold sequences using Bionano Solve (Bionano Genomics) with default parameters.

### Genetic mapping and pseudomolecule sequence construction

DNA was extracted from all individuals of the mapping populations and their parental lines using the sbeadex DNA extraction kit on the oKtopure system (LGC, United Kingdom). The extracted DNA was digested with PstI and MspI restriction endonucleases and subjected to ddRAD-Seq library preparation (Shirasawa *et al*. 2016). The resulting libraries were sequenced on a DNBSEQ G400 instrument (MGI Tech) to generate 100-bp paired-end reads. Alternatively, specific populations were genotyped using GRAS-Di (Enoki and Takeuchi 2018), and the sequence reads were processed as described below. After removing adapter sequences (AGATCGGAAGAGC for ddRAD-Seq or CTGTCTCTTATACACATCT for GRAS-Di) using fastx_clipper in the FASTX-Toolkit (version 0.0.14) and trimming low-quality reads (quality score < 10) using PRINSEQ (version 0.20.4) (Schmieder and Edwards 2011), high-quality reads were aligned to the hybrid scaffold sequences using Bowtie2 (version 2.3.5.1) (Langmead and Salzberg 2012). High-confidence biallelic SNPs were identified using the mpileup and call options of BCFtools (version 1.9) (Li 2011) and filtered using VCFtools (version 0.1.16) (Danecek *et al*. 2011) based on the following criteria: read depth ≥ 5; SNP quality = 999; minor allele frequency ≥ 0.2; and proportion of missing data < 20%. Linkage analysis of the SNPs was performed using Lep-Map3 (version 0.2) (Rastas 2017) to construct a genetic map. Finally, the hybrid scaffold sequences were anchored to the genetic map to establish chromosome-level pseudomolecule sequences using ALLMAPS (version 0.7.3) (Tang *et al*. 2015).

### Transcriptome analysis with long-and short-read sequencing

Total RNA was extracted from four tissues (male and female strobili, stems, and leaves) of C. obtusa and three tissues (shoots, stems, and leaves) of C. lanceolata using the FavorPrep Plant Total RNA Extraction Kit (Favorgen Biotech Corp, Ping Tung, Taiwan). Full-length RNA libraries for long-read cDNA sequencing were prepared using the Iso-Seq Express 2.0 Kit or the Kinnex Full-Length RNA Kit, and sequenced on the Sequel II or Revio systems (PacBio, Menlo Park, CA, USA). The resulting full-length cDNA sequences were processed using the Iso-Seq pipeline to generate non-redundant transcript sequences.

RNA libraries for short-read cDNA sequencing were prepared using the TruSeq Stranded mRNA Sample Preparation Kit (Illumina) and sequenced on a DNBSEQ G400 instrument (MGI Tech, Shenzhen, China). Adapter sequences and low-quality bases were trimmed as described above.

### Gene and repetitive sequence annotation

Protein-coding genes were predicted in three steps: 1) Transcript-based prediction was performed using GeneMarkS-T (Besemer and Borodovsky 2005) with full-length cDNA sequences generated from Iso-Seq or Kinnex; 2) *Ab initio* prediction was conducted using Helixer (Stiehler *et al*. 2021); and 3) BRAKER2 (Brůna *et al*. 2021) was used in conjunction with short-read RNA-Seq data to identify genes missed by the first two methods. Gene prediction completeness was evaluated using Benchmarking Universal Single-Copy Orthologs (BUSCO) with default parameters (Simão *et al*. 2015) and the gymnosperms_odb10 lineage dataset (Wu *et al*. 2023).

Repetitive sequences in the genome assemblies were identified using RepeatMasker (https://www.repeatmasker.org) with reference repeat sequences from Repbase and *de novo* repeat libraries constructed using RepeatModeler (https://www.repeatmasker.org).

### Comparative analysis of the genome structure

Pairwise collinear blocks were identified using MCScanX (Wang *et al*. 2012) with a match score threshold (-k) of 120 and an E-value cutoff of 10^-50^. The results were visualized using the synteny browser SynVisio (Bandi and Gutwin 2020).

### Organelle genome assembly and annotation

Chloroplast and mitochondrial genomes were assembled using HiFi reads obtained in our previous study (Shirasawa *et al*. 2024) via Oatk (version 1.0), with the syncmer coverage threshold (-c) set to 30 (Zhou *et al*. 2025). While Oatk successfully generated both chloroplast and mitochondrial sequences for *C. obtusa* and *C. lanceolata*, it did not automatically assemble the mitochondrial genome of *C. japonica*. Consequently, we extracted mitochondrial unitigs from the Oatk assembly graph—identified via HMM search—and manually constructed a contig using Bandage (version 0.9.0) (Wick *et al*. 2015).

Chloroplast genes (protein-coding, tRNA, and rRNA) were annotated using LiftOn (version 1.0.5) (Chao *et al*. 2025) based on alignments to published reference genomes: *C. obtusa* (GenBank accession PV469661.1), *C. japonica* (NC_010548.1), and *C. lanceolata* (NC_021437.1). Mitochondrial genes were similarly annotated using LiftOn, utilizing references from *Platycladus orientalis* (OL703044.1–OL703045.1), *Thuja sutchuenensis* (ON603305.1–ON603308.1), *Taxus chinensis* (NC_069591.1), *Taxus wallichiana* (NC_072607.1), and *Taxus cuspidata* (MN593023.1). Additionally, mitochondrial genes were predicted using PMGA (Li *et al*. 2025), a pipeline for plant mitochondrial genome annotation. All gene structures were manually curated and corrected, and final annotations were visualized using OGDRAW (version 1.3.1) (Greiner *et al*. 2019).

## Results

### Genome assembly and gene annotation for C. obtusa

In a previous study (Shirasawa *et al*. 2024), we established a draft genome assembly for *C. obtusa* containing primary contigs spanning 8.5 Gb (N50 = 1.2 Mb) and alternate contigs spanning 7.8 Gb (N50 = 496.6 kb). To extend sequence contiguity, the combined 16.3 Gb of primary and alternate sequences were scaffolded using 2,058.1 Gb of optical mapping data (7,538,216 molecules; N50 length = 289.7 kb). The resulting hybrid scaffold assembly comprised 848 sequences with a total length of 9.6 Gb and an N50 of 54.2 Mb.

Subsequently, four genetic maps of *C. obtusa* were constructed via linkage analysis of SNPs across four mapping populations. Each genetic map consisted of 11 linkage groups (Supplementary Table S1), corresponding to the basic chromosome number of *C. obtusa*. Based on these genetic maps, 378 hybrid scaffold sequences were anchored to the 11 linkage groups, establishing chromosome-scale pseudomolecule sequences spanning a total of 8.7 Gb (Table 1). The remaining 470 sequences (totaling 899.6 Mb) were unplaced.

**Table 1.** Statistics of the chromosome-scale genome assemblies for *C. obtusa*, *C. japonica*, and *C. lanceolata*.

| Chromosome | *C. obtusa* |  | *C. japonica* |  | *C. lanceolata* |  |
| --- | --- | --- | --- | --- | --- | --- |
|  | Length (bp) | % of length | Length (bp) | % of length | Length (bp) | % of length |
| Chr. 1 | 753,361,174 | 7.8 | 808,184,979 | 7.0 | 1,071,765,491 | 6.3 |
| Chr. 2 | 643,942,463 | 6.7 | 787,791,099 | 6.8 | 1,098,708,507 | 6.5 |
| Chr. 3 | 1,044,836,393 | 10.8 | 1,101,135,464 | 9.5 | 1,959,509,874 | 11.6 |
| Chr. 4 | 815,834,336 | 8.5 | 803,702,909 | 6.9 | 1,350,266,392 | 8 |
| Chr. 5 | 926,357,518 | 9.6 | 1,043,635,872 | 9.0 | 1,594,571,851 | 9.4 |
| Chr. 6 | 607,243,501 | 6.3 | 760,652,447 | 6.5 | 906,465,489 | 5.4 |
| Chr. 7 | 938,369,984 | 9.7 | 921,502,449 | 7.9 | 1,534,292,120 | 9.1 |
| Chr. 8 | 754,287,409 | 7.8 | 787,691,810 | 6.8 | 922,755,377 | 5.5 |
| Chr. 9 | 651,761,957 | 6.8 | 828,523,594 | 7.1 | 734,130,669 | 4.3 |
| Chr. 10 | 888,336,793 | 9.2 | 950,971,039 | 8.2 | 1,497,158,838 | 8.8 |
| Chr. 11 | 722,417,066 | 7.5 | 797,504,455 | 6.9 | 753,462,524 | 4.5 |
| **Subtotal** | **8,746,748,594** | **90.7** | **9,591,296,117** | **82.5** | **13,423,087,132** | **79.3** |
| Unplaced contigs | 899,622,983 | 9.3 | 2,035,395,917 | 17.5 | 3,497,085,311 | 20.7 |
| **Total** | **9,646,371,577** | **100.0** | **11,626,692,034** | **100.0** | **16,920,172,443** | **100** |

For gene prediction, transcriptome sequences were generated using both long-and short-read sequencing. Full-length cDNA sequences were clustered into 456,139 sequences containing 283,181 isoforms. From these, 27,384 non-redundant sequences were selected to predict 19,938 gene models, achieving a complete BUSCO score of 82.6%. In parallel, 1,728 million RNA-Seq reads from 96 samples were used to predict 41,266 gene models (complete BUSCO score: 61.1%). Furthermore, the *ab initio* method predicted 50,494 gene models (complete BUSCO score: 60.6%). Finally, we merged these three datasets to select a total of 54,034 non-redundant gene models (Table 2), achieving a final complete BUSCO score of 90.1% (Table 3).

**Table 2.** Summary of predicted genes in the genomes of *C. obtusa*, *C. japonica*, and *C. lanceolata*.

| Chromosome | *C. obtusa* |  | *C. japonica* |  | *C. lanceolata* |  |
| --- | --- | --- | --- | --- | --- | --- |
|  | #Genes | % of genes | #Genes | % of genes | #Genes | % of genes |
| Chr. 1 | 4,303 | 8.0 | 3,325 | 6.7 | 4,651 | 6.4 |
| Chr. 2 | 4,042 | 7.5 | 3,323 | 6.7 | 4,808 | 6.7 |
| Chr. 3 | 5,928 | 11.0 | 5,037 | 10.1 | 7,902 | 10.9 |
| Chr. 4 | 4,605 | 8.5 | 3,503 | 7.0 | 5,688 | 7.9 |
| Chr. 5 | 5,047 | 9.3 | 4,421 | 8.9 | 6,207 | 8.6 |
| Chr. 6 | 3,337 | 6.2 | 2,958 | 5.9 | 4,320 | 6 |
| Chr. 7 | 5,661 | 10.5 | 4,192 | 8.4 | 6,612 | 9.2 |
| Chr. 8 | 3,829 | 7.1 | 2,992 | 6.0 | 3,963 | 5.5 |
| Chr. 9 | 3,922 | 7.3 | 3,241 | 6.5 | 3,152 | 4.4 |
| Chr. 10 | 4,957 | 9.2 | 3,943 | 7.9 | 6,135 | 8.5 |
| Chr. 11 | 3,760 | 7.0 | 3,117 | 6.2 | 3,485 | 4.8 |
| **Subtotal** | **49,391** | **91.4** | **40,052** | **80.3** | **56,923** | **78.9** |
| Unplaced contigs | 4,643 | 8.6 | 9,847 | 19.7 | 15,246 | 21.1 |
| **Total** | **54,034** | **100.0** | **49,899** | **100.0** | **72,169** | **100** |

**Table 3.** Completeness assessment of gene predictions for *C. obtusa*, *C. japonica*, and *C. lanceolata*.

|  | *C. obtusa* | *C. japonica* | *C. lanceolata* |
| --- | --- | --- | --- |
| Complete | 90.1% | 90.5% | 93.8% |
| Single-copy complete | 77.5% | 76.0% | 59.6% |
| Duplicated complete | 12.6% | 14.5% | 34.2% |
| Fragmented | 0.7% | 0.9% | 0.6% |
| Missing | 9.2% | 8.6% | 5.6% |

### Genome assembly and gene annotation for C. japonica

The draft genome assembly for *C. japonica* comprised primary contigs spanning 9.2 Gb (N50 = 8.3 Mb) and alternate contigs spanning 8.7 Gb (N50 = 2.1 Mb) (Shirasawa *et al*. 2024). A total of 17.9 Gb of sequences was scaffolded using 2,151.5 Gb of optical mapping data (7,318,858 molecules; N50 length = 288.9 kb). The resulting hybrid scaffold assembly consisted of 805 sequences with a total length of 11.6 Gb and an N50 of 37.9 Mb.

Five genetic maps were constructed for *C. japonica* using five mapping populations. Each genetic map consisted of 11 linkage groups (Supplementary Table S2), corresponding to the basic chromosome number of *C. japonica*. Based on these maps, 294 hybrid scaffold sequences were anchored to the 11 linkage groups to establish chromosome-scale pseudomolecules spanning 9.6 Gb (Table 1). The remaining 511 sequences (2.0 Gb total) were unplaced.

For gene prediction, we used 75,348 isoforms derived from full-length cDNA sequences to predict 14,449 gene models (BUSCO completeness: 83.4%). Next, we utilized 34,731 unigene sequences from our previous study (Mishima *et al*. 2018) to predict 25,856 gene models (BUSCO completeness: 59.9%). Additionally, the *ab initio* method predicted 58,119 gene models (BUSCO completeness: 61.4%). These three datasets were merged to select a final set of 49,899 non-redundant gene models (Table 2), achieving a complete BUSCO score of 90.5% (Table 3).

### Genome assembly and gene annotation for C. lanceolata

The draft genome assembly for *C. lanceolata* contained primary contigs spanning 11.5 Gb (N50 = 11.7 Mb) and alternate contigs spanning 11.0 Gb (N50 = 2.9 Mb) (Shirasawa *et al*. 2024). A total of 22.5 Gb of sequence was scaffolded using 2,191.6 Gb of optical mapping data (5,368,583 molecules; N50 = 397.9 kb). The resulting hybrid scaffold assembly comprised 822 sequences with a total length of 16.9 Gb and an N50 of 31.8 Mb.

Two genetic maps for *C. lanceolata* were constructed using two distinct mapping populations. One genetic map consisted of 11 linkage groups (Supplementary Table S3), corresponding to the basic chromosome number of *C. lanceolata*. The second map consisted of 16 linkage groups (Supplementary Table S3); in six cases, pairs of linkage groups were assigned to a single chromosome sequence, while one chromosome was not represented by any linkage group. Ultimately, 298 hybrid scaffold sequences were anchored to the 11 linkage groups to establish chromosome-scale pseudomolecule sequences spanning a total of 13.4 Gb (Table 1).

The remaining 524 sequences (3.5 Gb total) were unplaced.

For gene prediction, we utilized 1,088,667 non-redundant full-length cDNA sequences to predict 35,735 gene models (complete BUSCO score: 90.5%). Next, 113.2 million RNA-Seq reads from stem, root, and leaf samples (NCBI SRA accession numbers SRR5282557–SRR5282559) were used to predict 47,922 gene models (complete BUSCO score: 62.0%). Additionally, the *ab initio* method predicted 45,934 gene models (complete BUSCO score: 40.6%). These three datasets were merged to select a total of 72,169 non-redundant gene models (Table 2), achieving a final complete BUSCO score of 93.8% (Table 3).

### Repetitive sequence analysis

Repetitive sequences account for 77.5%, 80.5%, and 80.9% of the genomes of *C. obtusa*, *C. japonica*, and *C. lanceolata*, respectively. LTR retrotransposons were the most abundant class of repetitive sequences (48.5–53.6%), followed by unclassified repeats (14.3–21.6%), DNA transposons (5.2–6.8%), and LINEs (3.3–4.9%) (Table 4).

**Table 4.** Classification and abundance of repetitive sequences in the genomes of *C. obtusa*, *C. japonica*, and *C. lanceolata*.

| Repeat type | *C. obtusa* |  |  | *C. japonica* |  |  | *C. lanceolata* |  |  |
| --- | --- | --- | --- | --- | --- | --- | --- | --- | --- |
|  | #Elements | Length (bp) | % | #Elements | Length (bp) | % | #Elements | Length (bp) | % |
| SINEs | 11,430 | 1,313,246 | 0.0 | 12,828 | 2,413,789 | 0.0 | 50,707 | 15,072,777 | 0.1 |
| LINEs | 536,344 | 356,067,530 | 3.7 | 647,369 | 568,704,286 | 4.9 | 793,194 | 558,315,070 | 3.3 |
| LTR elements | 2,900,360 | 4,680,905,729 | 48.5 | 3,606,721 | 6,235,301,883 | 53.6 | 6,005,684 | 8,319,335,602 | 49.2 |
| DNA transposons | 738,498 | 505,285,736 | 5.2 | 1,137,352 | 785,356,255 | 6.8 | 1,277,378 | 903,211,596 | 5.3 |
| Small RNA | 9,934 | 4,679,246 | 0.1 | 23,970 | 9,465,882 | 0.1 | 68,538 | 20,640,162 | 0.1 |
| Satellites | 29,335 | 26,779,510 | 0.3 | 19,979 | 10,042,049 | 0.1 | 75,861 | 45,335,388 | 0.3 |
| Simple repeats | 764,656 | 55,374,637 | 0.6 | 902,078 | 50,675,895 | 0.4 | 980,278 | 58,458,100 | 0.4 |
| Low complexity | 125,181 | 8,185,083 | 0.1 | 144,624 | 8,419,661 | 0.1 | 166,010 | 10,190,191 | 0.1 |
| Unclassified | 3,210,805 | 1,807,553,850 | 18.7 | 3,289,375 | 1,666,957,226 | 14.3 | 8,058,531 | 3,655,997,225 | 21.6 |

### Comparative analysis of the three coniferous genomes

The chromosome-scale genome sequences of the three species were compared pairwise (Figure 1A). This comparison revealed that while the structures of eight chromosomes (Chr 1, 3, 4, 5, 7, 8, 10, and 11) were well conserved across all three species, three chromosomes (Chr 2, 6, and 9) underwent significant rearrangements (Figure 1B).

**Figure 1.**
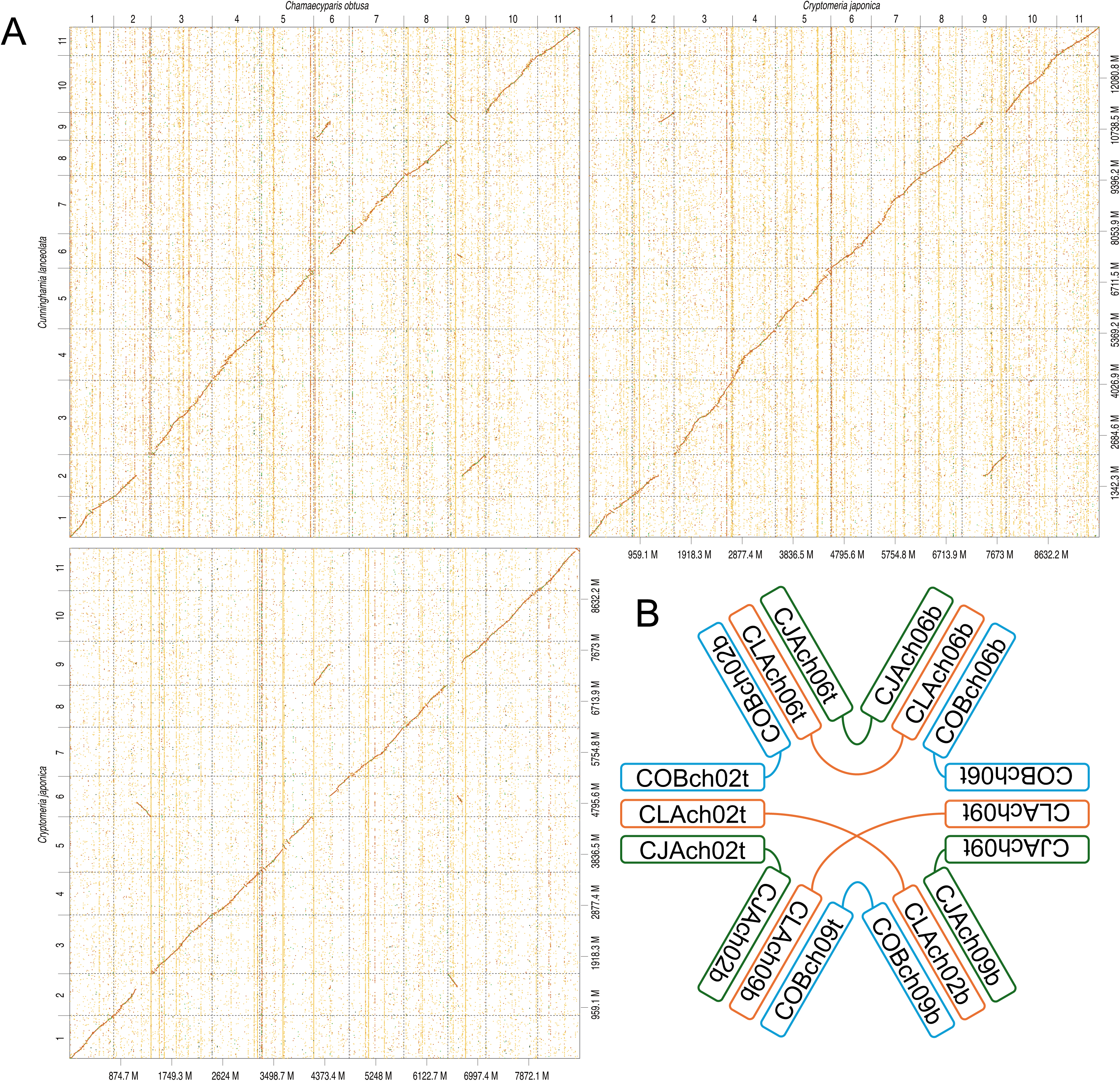
Comparative analysis of genome sequences and structures of *C. obtusa*, *C. japonica*, and *C. lanceolata*. Chromosome numbers are indicated on the top (x-axis) and left (y-axis). Genome sizes (Mb) are shown on the bottom (x-axis) and right (y-axis).

In *C. obtusa* chromosome 2, the top region (COBch02t) was conserved across all three species. However, the bottom region (COBch02b) was inverted and translocated to the top of chromosome 6 in both *C. japonica* (CJAch06t) and *C. lanceolata* (CLAch06t). Concurrently, regarding *C. obtusa* chromosome 6, the bottom region was conserved across species, while the top region (COBch06t) was translocated to the top of chromosome 9 in *C. japonica* (CJAch09t) and *C. lanceolata* (CLAch09t).

The structure of *C. obtusa* chromosome 9 was more complex. The top region (COBch09t) was inverted and translocated to the bottom of *C. japonica* chromosome 2 (CJAch02b) and the bottom of *C. lanceolata* chromosome 9 (CLAch09b). Conversely, the bottom region (COBch09b) was translocated to the bottom of *C. japonica* chromosome 9 (CJAch09b) and the bottom of *C. lanceolata* chromosome 2 (CLAch02b). These patterns indicate a reciprocal translocation event involving the bottom regions of chromosomes 2 and 9 between *C. japonica* and *C. lanceolata*.

Synteny analysis (Figure 2A) identified 216 pairwise collinear blocks (17,449 gene pairs) between *C. obtusa* and *C. japonica*. Similarly, 271 blocks (16,918 gene pairs) were identified between *C. japonica* and *C. lanceolata*, and 254 blocks (16,022 gene pairs) were identified between *C. lanceolata* and *C. obtusa*. Finally, the analysis detected tandem gene duplications (Figure 2B), revealing 23 blocks (525 gene pairs) in *C. obtusa*, 37 blocks (924 gene pairs) in *C. japonica*, and a notably higher frequency of 89 blocks (3,149 gene pairs) in *C. lanceolata*.

**Figure 2.**
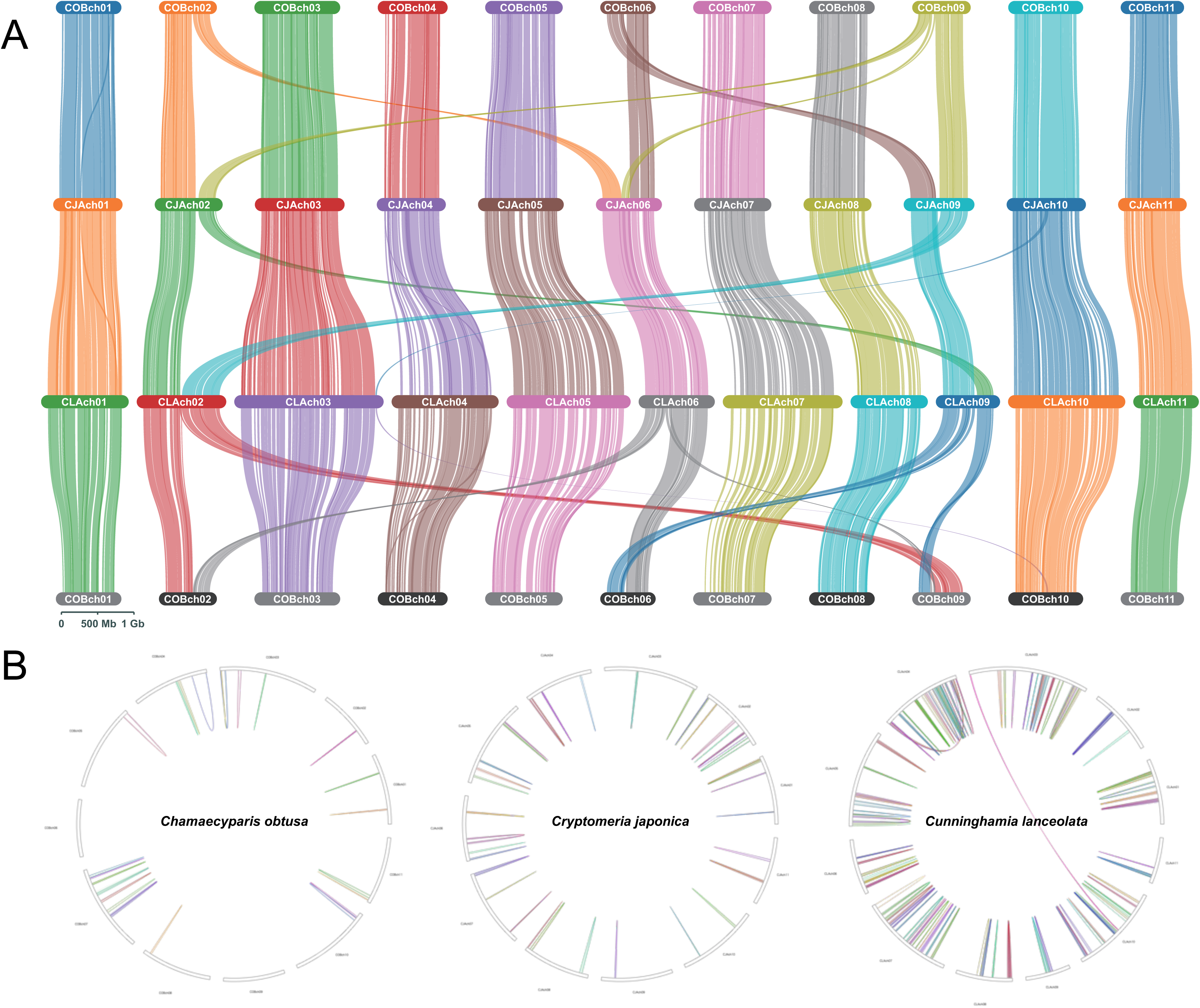
Synteny analysis of *C. obtusa*, *C. japonica*, and *C. lanceolata*. (A) Lines connect pairwise collinear blocks between the genomes of *C. obtusa*, *C. japonica*, and *C. lanceolata*. (B) Lines connect collinear blocks within each genome to visualize tandem duplications.

### Organelle genome assemblies and gene annotations

The chloroplast genomes of *C. obtusa*, *C. japonica*, and *C. lanceolata* were assembled as single circular sequences with lengths of 127,726 bp, 131,741 bp, and 135,331 bp, respectively (Figure 3A). The *C. obtusa* chloroplast genome contained 120 genes (83 protein-coding, 33 tRNA, and 4 rRNA). In *C. japonica*, 118 genes were annotated (82 protein-coding, 32 tRNA, and 4 rRNA), while in *C. lanceolata*, 120 genes were annotated (81 protein-coding, 35 tRNA, and 4 rRNA).

**Figure 3.**
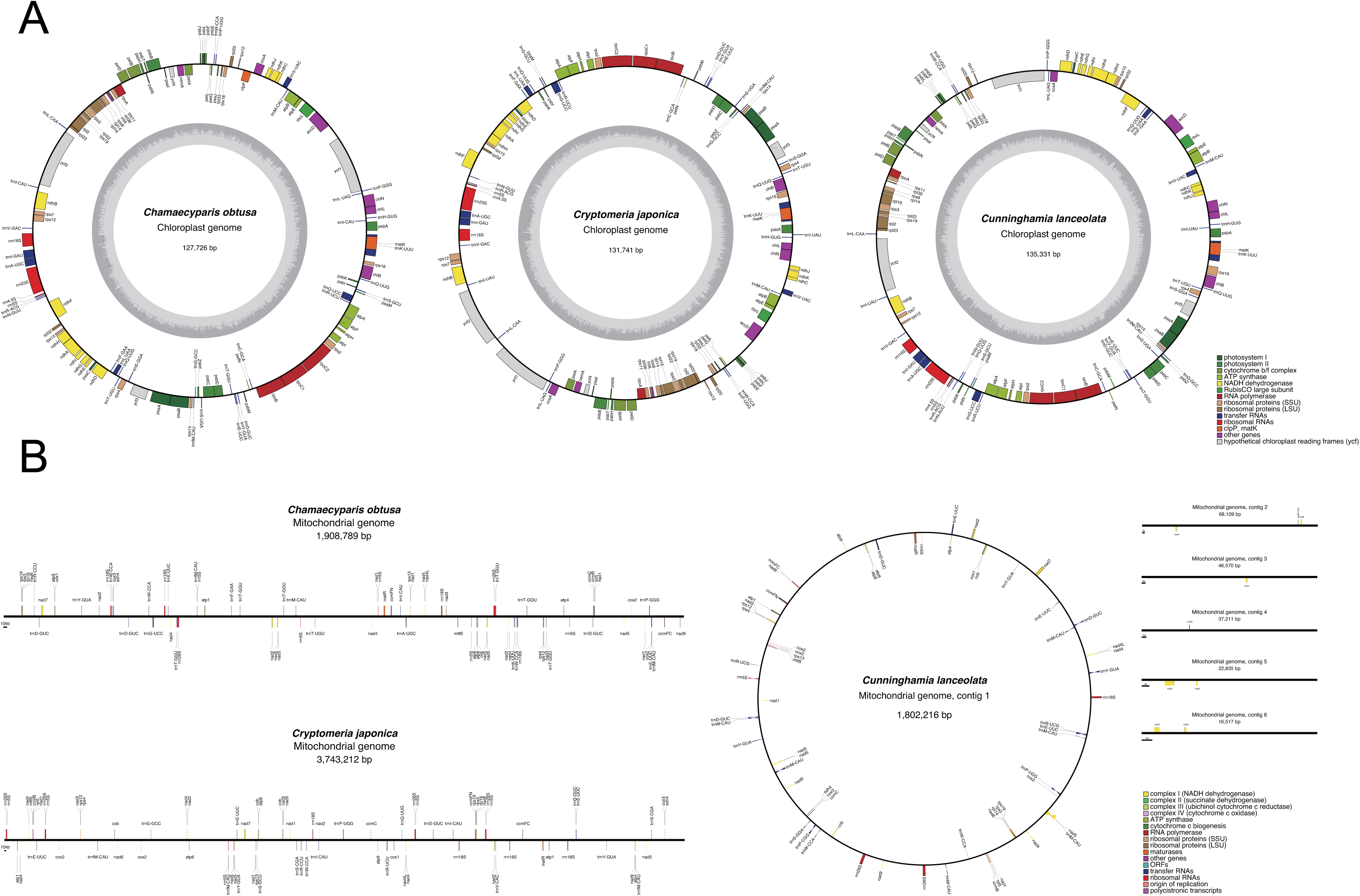
Organelle genomes of *C. obtusa*, *C. japonica*, and *C. lanceolata*. Chloroplast and mitochondrial genome structures are represented by solid lines. Boxes on the genome tracks indicate genes. A key to the gene functions is provided in the bottom-right corner.

The mitochondrial genome of *C. obtusa* was assembled as a single linear sequence of 1,908,789 bp, and that of *C. japonica* as a single linear sequence of 3,743,212 bp (Figure 3B). In contrast, the mitochondrial genome of *C. lanceolata* consisted of six sequences with a total length of 1,993,458 bp, comprising one circular sequence and five linear sequences (Figure 3B). Gene annotation identified 70 genes in the *C. obtusa* mitochondrial genome (33 protein-coding, 26 tRNA, 11 rRNA); 71 genes in *C. japonica* (36 protein-coding, 22 tRNA, 13 rRNA); and 65 genes in *C. lanceolata* (36 protein-coding, 23 tRNA, 6 rRNA).

## Discussion

We present chromosome-scale genome assemblies for three species of the Cupressaceae: *C. obtusa* (Japanese cypress), *C. japonica* (Japanese cedar), and *C. lanceolata* (Chinese fir) (Table 1). Consistent with previous genome size estimates (Shirasawa *et al*. 2024), the resulting assembly length of *C. lanceolata* (13.4 Gb) was significantly larger than those of *C. obtusa* (8.7 Gb) and *C. japonica* (9.6 Gb). Since the proportion of repetitive sequences remained approximately 80% across all three genomes (Table 4), the larger genome size of *C. lanceolata* appears to be driven by the multiplication of genomic segments containing both repetitive and non-repetitive sequences, rather than solely by a burst of transposon activity. This hypothesis is supported by the higher number of predicted genes in *C. lanceolata* (72,169) compared to *C. obtusa* (54,034) and *C. japonica* (49,899) (Table 2), as well as its elevated duplicated BUSCO score (34.2%) relative to *C. obtusa* (12.6%) and *C. japonica* (14.5%) (Table 3). These results suggest that frequent tandem gene duplications (Figure 2B) have contributed to the expansion of both genome size and gene content in *C. lanceolata*.

Comparative genome analysis revealed structural rearrangements in three chromosomes (2, 6, and 9) between *C. obtusa* and *C. japonica*—a pattern consistent with a previous study (Dogan *et al*. 2024)— as well as between *C. obtusa* and *C. lanceolata*. In contrast, rearrangements were observed in only two chromosomes (2 and 9) between *C. japonica* and *C. lanceolata* (Figure 1). Based solely on the number of chromosomal rearrangements, one might infer a closer relationship between *C. japonica* and *C. lanceolata*. However, phylogenetic studies (Liu *et al*. 2022) indicate that *Cunninghamia* diverged earlier from the common ancestor of *Chamaecyparis* and *Cryptomeria*. The chromosomal rearrangements and tandem gene duplications identified in this study (Figures 1 and 2) provide new insights into the complex processes of evolution and speciation in these coniferous trees.

In conclusion, this study provides essential genomic resources, including chromosome-scale assemblies, gene annotations, and genetic maps, for *C. obtusa*, *C. japonica*, and *C. lanceolata*. Although one chromosome-scale assembly for *C. japonica* was recently published (Fujino *et al*. 2024), few other genomic resources have been publicly available for these species. The availability of multiple genome assemblies, facilitating pan-genome analysis, is crucial for understanding the genetic mechanisms underlying species diversity (Aoyagi Blue *et al*. 2025). The resources presented here, combined with existing data, will facilitate further research into the evolution and speciation of these conifers and accelerate the identification of key genes for breeding programs.

## Acknowledgements

We thank Y. Kishida, C. Minami, K. Ozawa, H. Tsuruoka, and A. Watanabe (Kazusa DNA Research Institute) for technical assistance.

## Funding

This study was supported in part by the MAFF commissioned project study on “Development of efficient breeding technique aiming at forestry trees with superior carbon storage capacity” (Grant Number JPJ009841), JSPS KAKENHI (22H05172 and 22H05181), and the Kazusa DNA Research Institute Foundation.

## Data availability

The genome assembly files and gene annotation files for the three species are available in Kazusa Genome Atlas (https://genome.kazusa.or.jp).

## Conflicts of interest

All authors declare no competing interests.

## Notes

### Competing Interest Statement

The authors have declared no competing interest.

